# Reelin Mediates Hippocampal Cajal-Retzius Cell Positioning and Infrapyramidal Blade Morphogenesis

**DOI:** 10.1101/543074

**Authors:** Seungshin Ha, Prem P. Tripathi, Ray A. Daza, Robert F. Hevner, David R. Beier

**Affiliations:** Center for Developmental Biology and Regenerative Medicine, Seattle Children’s Research Institute, Seattle, WA 98101; Division of Genetic Medicine, Department of Pediatrics, University of Washington School of Medicine, Seattle, WA 98195; Center for Integrative Brain Research, Seattle Children’s Research Institute, Seattle, WA 98101; Department of Neurological Surgery, University of Washington School of Medicine, Seattle, WA 98195

**Keywords:** reelin protein, hippocampus, dentate gyrus, postnatal development, migration, neurogenesis

## Abstract

We have previously described hypomorphic *reelin* (*Reln*) mutant mice, *Reln*^*CTRdel*^, in which the morphology of the dentate gyrus is distinct from that seen in *reeler* mice. In the *Reln*^*CTRdel*^ mutant the infrapyramidal blade of the dentate gyrus fails to extend, while the suprapyramidal blade forms with a relatively compact granule neuron layer. The distribution of Cajal-Retzius cells in the dentate gyrus was aberrant; Cajal-Retzius neurons were increased in the suprapyramidal blade, but were greatly reduced along the subpial surface of the prospective infrapyramidal blade. We also observed multiple abnormalities of the fimbriodentate junction. Firstly, progenitor cells were distributed abnormally; the “neurogenic cluster” at the fimbriodentate junction was absent, lacking the normal accumulation of Tbr2-positive intermediate progenitors. However, the number of dividing cells in the dentate gyrus was not generally decreased. Secondly, a defect of secondary glial scaffold formation, limited to the infrapyramidal blade, was observed. The densely radiating glial fibers characteristic of the normal fimbriodentate junction were absent in mutants. These fibers might be required for migration of progenitors, which may account for the failure of neurogenic cluster formation. These findings suggest the importance of the secondary scaffold and neurogenic cluster of the fimbriodentate junction in morphogenesis of the mammalian dentate gyrus. Our study provides direct genetic evidence showing that normal RELN function is required for Cajal-Retzius cell positioning in the dentate gyrus, and for formation of the fimbriodentate junction to promote infrapyramidal blade extension.

## INTRODUCTION

Morphogenesis of the dentate gyrus in rodents begins during mid-gestation, and the key events for infrapyramidal blade (IPB) elongation occur during the first postnatal week (Altman and Bayer, 1990a; Altman and Bayer, 1990b; Li and Pleasure, 2005; Forster et al., 2006; Hodge et al., 2013; Hayashi et al., 2015). The granule neuron progenitors are born in the dentate neuroepithelium during the late embryonic and early postnatal period. The Cajal-Retzius cells expressing RELN migrate into the invagination of the hippocampal fissure above the suprapyramidal blade (SPB). The progenitor cells migrate along the dentate migratory stream (parallel and adjacent to the Cajal-Retzius cell migration) to the developing dentate gyrus, toward the sources of RELN in the hippocampal fissure. At birth, the SPB of the granule cell layer has already formed along the hippocampal fissure, but the IPB has only just begun to grow along the medial subpial surface. As the IPB extends, the Cajal-Retzius cells also migrate and disperse along the subpial surface of the prospective IPB.

The secondary glial scaffold forms initially in the SPB, as glial fibers radiate from the medial edge of the subpial zone (fimbriodentate junction) across the dentate hilus to the hippocampal sulcus. Progenitors from the dentate migration stream pause and accumulate at the fimbriodentate junction, where a fork in the road is encountered (as previously described in Figure 1C of Hodge et al., 2013). From the fimbriodentate junction, progenitors further migrate into the hilus (transhilar stream) or the dentate molecular layer (subpial stream). Tangential growth (elongation) of the infrapyramidal blade depends on the coordinated migration and accretion of new radial glial scaffold fibers, Cajal-Retzius neurons, granule neurons, and neural progenitors, including neural stem cells with morphological and molecular properties similar to radial glia. By the end of the first postnatal week, the radial fibers and outer neuronal shell of the infrapyramidal blade are formed. Granule neurons are thereafter produced from several postnatal niches, including the hilus, subpial neurogenic zone, fimbriodentate junction, and subgranular zone (Li et al., 2009; Nicola et al., 2015).

*Reln* deficiency causes a severe malformation of the dentate gyrus. The dentate granule neurons do not form a layered structure, but are loosely scattered in the hilus (Stanfield and Cowan, 1979a). The *reeler* phenotype suggested that RELN acts as a positional cue during dentate gyrus development (Frotscher et al., 2003; Forster et al., 2006; Zhao and Frotscher, 2010). When *reeler* and wild-type hippocampal slices are co-cultured next to each other, the dentate granule cell layer lamination is rescued by migration of neurons and glia toward the source of RELN (Zhao et al., 2004). *Reeler* does not develop the secondary glial scaffold at all, and the lack of this structure has been considered as the main cause of dentate gyrus malformation (Forster et al., 2002; Frotscher et al., 2003; Weiss et al., 2003). RELN receptor (*Vldlr or Apoer2*) deficient mice also have mild defects in the secondary glial scaffold, and develop a less compact granule cell layer (Weiss et al., 2003; Brunne et al., 2013). The dentate gyrus malformations seen in *reeler* or RELN receptor deficient mice are not specific to the IPB. On the other hand, IPB malformation has been associated with abnormal RELN expression (Yang et al., 2000; Tian et al., 2012; Hodge et al., 2013). However, in these cases the requirement of RELN for IPB development was still inconclusive due to the lack of direct genetic evidence proving causality; that is, *reeler* mice appear to develop the IPB (although it is highly disorganized) and a laminated outer molecular layer (Deller et al., 1999).

In a mutagenesis screen of ENU-treated mice, we identified a hypomorphic allele of *Reln, Reln*^*CTRdel*^ (Ha et al., 2015). This mutant carries a short C-terminal deletion of RELN protein. Studies using multiple mutant mice have revealed that C-terminal region domain (CTR) is important for secretion, proteolytic cleavage, receptor binding, activation of downstream signaling, resulting in abnormalities in brain morphogenesis and behavior (D’Arcangelo et al., 1997; de Bergeyck et al., 1997; Nakano et al., 2007; Kohno et al., 2015; Nakamura et al., 2016; Sakai et al., 2016). Using genetic and biochemical analyses, we have demonstrated that the C-terminal deletion resulted in differential binding of RELN to its receptors; truncated RELN fails to bind to VLDLR but binds normally to APOER2 (Ha et al., 2017).

The mutation appears to specifically disrupt IPB extension, as the mutant develops a compact SPB with only subtle disorganization (Ha et al., 2017). This is a unique phenotype of *Reln*^*CTRdel*^, as knock-in mice without CTR are able to develop IPB (Kohno et al., 2015; Sakai et al., 2016). To further characterize the IPB defect in the mutant, we examined several aspects of dentate gyrus development during the first postnatal week, focusing on those that have been found to be altered in *reeler* mice. We found that *Reln*^*CTRdel*^ mutants show abnormal distribution of the Cajal-Retzius cells, which has not been previously reported in *reeler* mice. Subsequent defects in glia scaffold development and neurogenesis at the fimbriodentate junction arising at early postnatal days led to more pronounced malformations in later stages, eventually resulting in truncation of the IPB.

## MATERIALS AND METHODS

### Mice

Generation and characterization of *Reln*^*CTRdel*^ mice (RRID: MGI:5505412) was previously described (Ha et al., 2015; Ha et al., 2017). The mutant mice have a mixed background, carrying A/J alleles in proximal chromosome 5 ∼28Mb region flanked by microsatellite markers D5Mit193 and D5Mit386. The mice analyzed were outcrossed at least 8 generations to C57BL6/N (Taconic). Mice of both sexes were analyzed. Animals were maintained in accordance with guidelines of National Institutes of Health and the Seattle Children’s Hospital Institutional Animal Care and Use Committee.

### Immunohistochemistry and histology

The brains were fixed by performing transcardial perfusion and then by incubating overnight in 4% paraformaldehyde in phosphate-buffered saline (PBS). Fixed brains were incubated overnight in 30% sucrose at 4 degree for cryoprotection, embedded in OCT media, and cryosectioned at 10 μm thickness. If necessary, antigen enhancement procedure was performed by boiling briefly in 10 mM sodium citrate (pH 6.0), followed by washing three times with PBS for 10 min each. For BrdU immunostaining, the sections were incubated in 2N hydrochloric acid at 37 degree for 1 hour, followed by washing six times with PBS for 10 min each. Blocking was performed for 30 min at room temperature by incubating in PBS-based blocking solution containing 5% normal goat serum, 0.3% Triton-X100, 2% bovine serum albumin. The sections were incubated with the primary antibodies (Table 1) overnight at 4 degree and washed three times with PBS for 10 min each. The secondary antibody incubation was done for 2 hours at room temperature. The secondary antibodies used are Alexa Fluor 488 or 568 goat anti-rabbit, anti-mouse, or anti-rat IgG (Life Technologies, 1:600). The sections were washed three times with PBS for 10 min each, stained using Hoechst (Life Technologies) following the manufacturer’s instruction, washed with PBS again before mounting.

**Table 1.** List of Primary Antibodies

List of Primary Antibodies
| Antigen | Description of immunogen | Source, host species, clonality, catalog No., clone No., RRID | Stock concentration, dilution used |
| --- | --- | --- | --- |
| Reelin | Recombinant reelin amino acids 164-496 | Millipore, mouse, monoclonal, Cat# MAB5364, clone G10, RRID:AB_2179313 | 1mg/mL, 1:1000 |
| p73 | Recombinant human p73α amino acids 1-80 | Santa Cruz, rabbit, polyclonal, Cat# sc-7957, clone H-79, RRID:AB_2207314 | 200μg/mL, 1:200 |
| BrdU | BrdU | Accurate, rat, monoclonal, Cat# OBT0030, clone BU1/75 (ICR1), RRID:AB_2313756 | 0.5mg/ml, 1:300 |
| Ki67 | Synthetic peptide derived from within amino acids 2300-2400 of human Ki67 | Vector Lab, rabbit, monoclonal, Cat# VP-Rm04, clone SP6, RRID:AB_2336545 | Unknown, 1:300 |
| Tbr2 | Mouse Tbr2/Eomes | eBioscience, rat, monoclonal, Cat# 14-4875-82, RRID:AB_11042577 | 0.5mg/mL, 1:300 |
| GFAP | GFAP isolated from cow spinal cord | Dako, rabbit, polyclonal, Cat# Z0334, RRID:AB_10013382 | 2.9mg/mL, 1:500 |
| BLBP | Synthetic peptide conjugated to KLH derived from within amino acids 1-100 of mouse BLBP | Abcam, rabbit, polyclonal, Cat# ab32423, RRID:AB_880078 | Unknown, 1:1000 |
| NeuroD | Peptide mapping at the N-terminus of mouse NeuroD | Santa Cruz, goat, polyclonal, Cat# sc-1084, clone N-19, RRID:AB_630922 | 100μg/ml, 1:400 |
| Prox1 | Synthetic peptide EIFKSPNCLQELLHE, corresponding to amino acids 723-737 of mouse Prox1 | Abcam, rabbit, polyclonal, Cat# ab37128, RRID:AB_882189 | Unknown, 1:1000 |

Nissl staining was done on frozen sections using Thionin solution (FD NeuroTechnologies) following the manufacturer’s instruction. Briefly, the sections were rehydrated, incubated in Thionin solution for 10 min and then in acidic alcohol for 2 min, dehydrated in ethanol and xylene, and mounted using Permount (Fisher Scientific).

### Antibody characterization

All antibodies used were purchased from commercial sources. The primary antibodies are listed in Table 1. All antibodies were selected based on the previously published characterization and utility. Anti-reelin clone G10 was developed and originally verified using Western blotting, immunoprecipitation, and immunohistochemistry (de Bergeyck et al., 1998; Pesold et al., 1998), and the clone G10 from Millipore (Cat# MAB5364, RRID: AB_2179313) was previously verified using Western blotting and immunohistochemistry in our hands (Ha et al., 2017). Anti-p73 (Santa Cruz, Cat# sc-7957, RRID: AB_2207314) was previously used for immunohistochemistry to label Cajal-Retzius cells (Gu et al., 2011), and also validated in our hands previously (Hodge et al., 2013). The use of antibodies for proliferative markers, anti-BrdU (Accurate, Cat# OBT0030, clone BU1/75-ICR1, RRID:AB_2313756) and anti-Ki67 (Vector Lab, Cat# VP-Rm04, clone SP6, RRID: AB_2336545) has been described previously (Hodge et al., 2012). GFAP (Dako, Cat# Z0334, RRID: AB_10013382) was previously used for immunohistochemistry to label glial fibers (Brunne et al., 2010; Brunne et al., 2013), and previously validated in our hands (Hodge et al., 2012). BLBP (Abcam, Cat# ab32423, RRID: AB_880078) was used to label radial glial scaffold (Gu et al., 2011), and previously validated in our hands (Hodge et al., 2012). Markers for granule neuron precursors, NeuroD (Santa Cruz, Cat# sc-1084, clone N-19, RRID: AB_630922) and Prox1 (Abcam, Cat# ab37128, RRID: AB_882189), were also previously used (Hodge et al., 2008; Pedroni et al., 2014). Staining results for all these tissue markers were replicates of the previously published distribution patterns and localization.

### BrdU incorporation assay

The pups were intraperitoneally injected with BrdU (50 mg/kg) at P3 and the brains were collected after 24 hours. Total 2 wild-type mice and 4 homozygote mutant mice were analyzed. The sections were double-labeled using anti-BrdU and anti-Ki67 antibodies. Cell counting was performed using ImageJ software (RRID:SCR_003070).

### Statistical analysis

Data are graphically expressed as boxplots indicating the median ± interquartile range (IQR) and whiskers indicating minimum and maximum. All data points are shown in the graph. 1-3 sections (replicates) per mice were analyzed, and the averaged counts from each mouse were used for statistical analysis. For statistical analysis, nonparametric Mann-Whitney test was used, and significance was considered when p <0.05 (two-tailed). The wild-type and mutant counts were treated as independent (unpaired) measurements. The statistical analysis was performed using GraphPad Prism7 (RRID: SCR_002798). Sample size calculation was performed using PS power and sample size program (http://biostat.mc.vanderbilt.edu/wiki/Main/PowerSampleSize).

## RESULTS

### Cajal-Retzius cells are absent in the IPB

In wild-type mice, RELN-expressing cells were found in the hippocampal fissure and the dentate crest at P0 (Ha et al., 2017), and in the subpial surface below the IPB at P7 (Figure 1A). The mutant had a relatively shorter region with RELN-expressing cells in the anterior IPB, which appeared truncated (Ha et al., 2017). Lack of RELN-expressing cells was strikingly obvious in the posterior sections, in which the IPB is completely absent (Figure 1A). This result was confirmed by examining the distribution of cells expressing the Cajal-Retzius cell marker p73 (Figure 1B). The numbers of cells expressing p73 appeared greater in the mutant SPB (Figure 1B). These results suggested that truncation of the IPB is due to an abnormality of Cajal-Retzius cell positioning.

**Figure 1.**
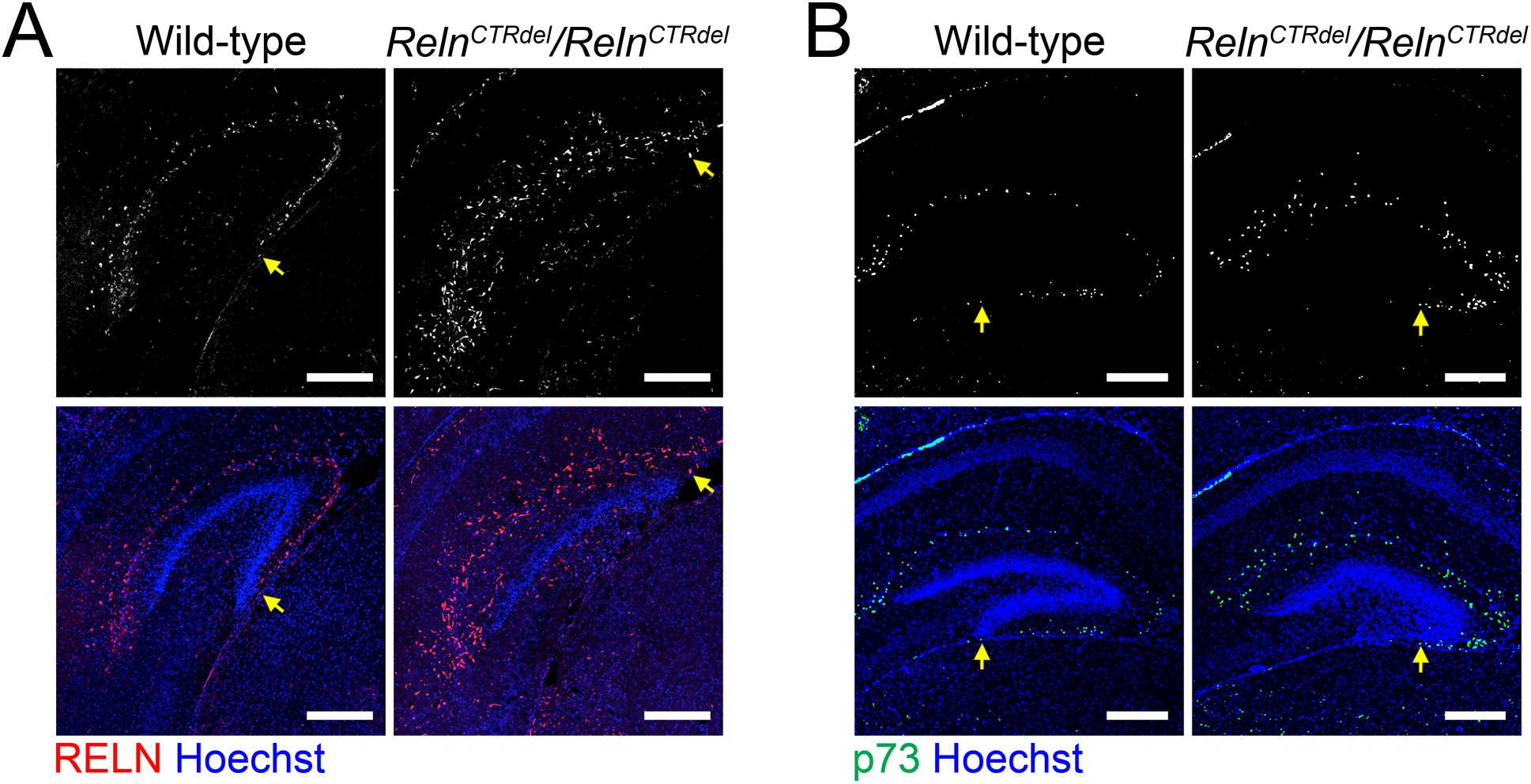
Lack of the Cajal-Retzius cells in the infrapyramidal blade. **A:** The infrapyramidal blade (IPB) of dentate gyrus does not develop in *Reln*^*CTRdel*^, and this coincides with the absence of the RELN-positive cells (gray or red) in the prospective infrapyramidal blade. P7 brains were immunostained with anti-RELN antibodies (n=2 each). **B:** Distribution of p73-expressing cells (gray or green) in the dentate gyrus at P7 (n=2 each). The number of p73-expressing cells is increased in the suprapyramidal blade (SPB). Nuclei were stained using Hoechst (blue). Scale bars, 250 μm. Arrows indicate the edge of Cajal-Retzius cell-containing regions, which are near the fimbrodentate junction (in the wild type) or the tip of truncated IPB (in the mutants).

### Neurogenic cluster does not form at the fimbriodentate junction

A remarkable aspect of the mutant phenotype is that the IPB is severely truncated, despite the fact that the mutant develops a granule cell layer in the SPB. This suggested that there might be a neurogenesis defect. The SPB of the dentate gyrus is established around birth in a primordial form of granule cell layer, followed by the formation of the IPB during the first postnatal week (Altman and Bayer, 1990a). We selected the middle time point to examine cell division, when the IPB formation is actively in progress. P3 pups were injected with BrdU and analyzed after 24 hours at P4, then immunostained with anti-BrdU and anti-Ki67 antibodies (Figure 2).

**Figure 2.**
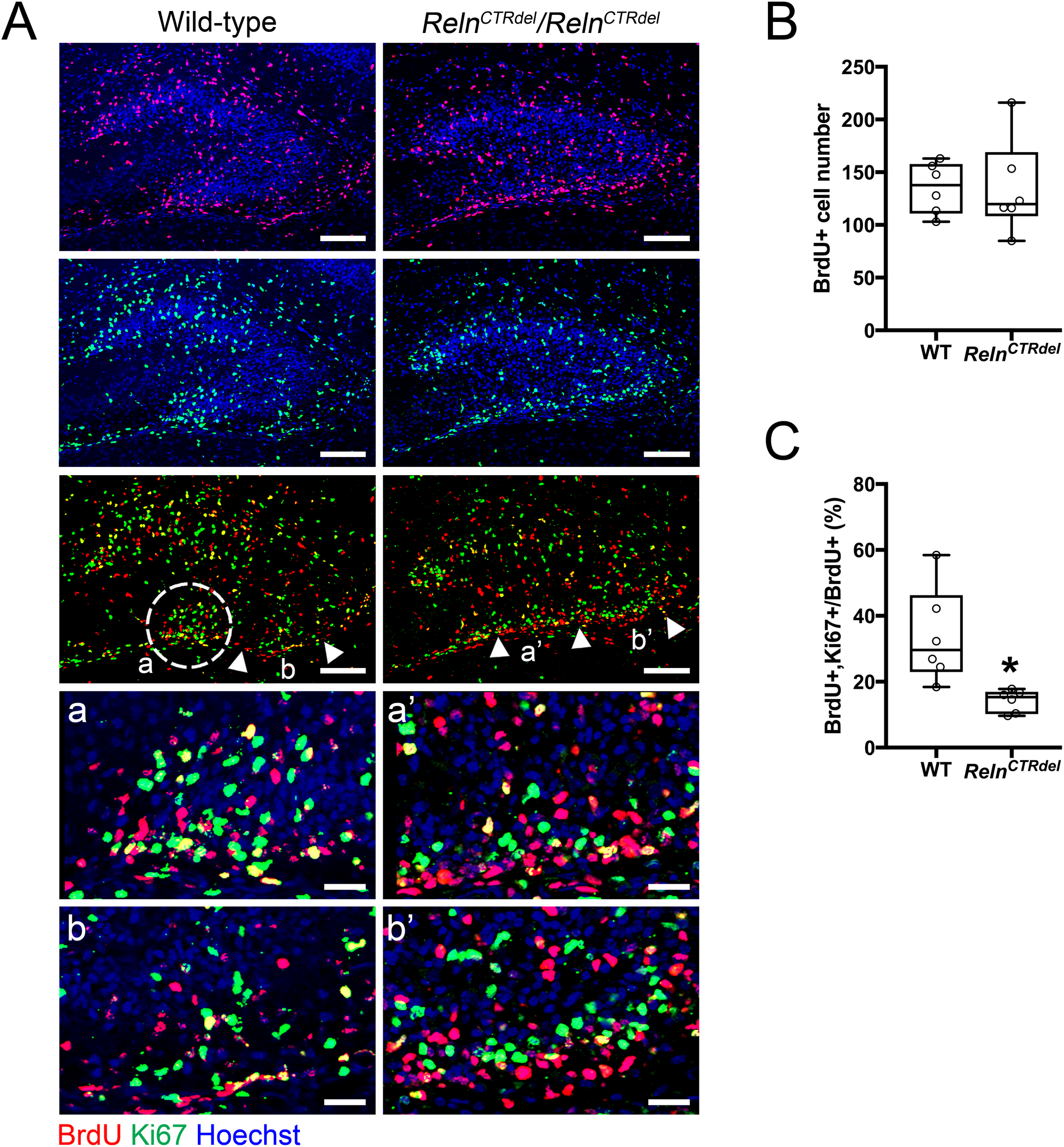
Absence of the neurogenic cluster at the fimbriodentate junction. **A:** Distribution of the BrdU (red) or Ki67-positive cells (green) in the dentate gyrus at P4, after 24 hours from BrdU injection. A cluster of dividing cells is present at the fimbriodentate junction of the wild-type (white circle, a). A *Reln*^*CTRdel*^ mutant does not have this cluster (a’). Compared with a corresponding region of the wild type (b), abnormal accumulation of the dividing cells are apparent in the subpial surface of *Reln*^*CTRdel*^ (b’). Enlarged images of marked regions (a, a’, b, b’) are shown below. In the mutant, Ki67-positive cells are closely located on top of BrdU-positive cells (a’ and b’). Scale bars, 100 μm. Scale bars in the enlarged images, 25 μm. Nuclei were stained using Hoechst (blue). **B:** The number of total BrdU-positive cells is not significantly different (p=0.8182; boxplot, median ± IQR; whiskers, min and max). **C:** The percentage of BrdU/Ki67 double-positive cells was reduced (p=0.0022; boxplot, median ± IQR; whiskers, min and max). This represents a population of cells that reentered the cell cycle.

The mutant displayed a markedly abnormal spatial distribution of dividing cells in the IPB region. In wild-type mice, a cluster of BrdU-positive cells and Ki67-positive cells was localized at the tip of growing granule cell layer (Figure 2A, dashed circle). In contrast, the mutants did not show a cluster of progenitors at the fimbriodentate junction; instead, dividing cells were dispersed along the subpial surface. Ki67-positive cells (dividing at the time of harvest) were localized within a thin layer above a layer of BrdU-positive cells (dividing at the time of injection) along the subpial zone. This result led us to speculate that these progenitor cells migrated inappropriately, due to defective guidance cues in mutant mice, especially at the fimbriodentate junction. Moreover, these observations suggested that the simple presence of progenitor cells near the prospective IPB is not sufficient for its formation, and that abnormal organization of the neurogenic niche contributes to the defective development of the IPB.

Of note, the total numbers of BrdU positive cells in the entire region of the dentate gyrus were not significantly different between wild-type and mutant (Figure 2B, p=0.8182, Mann-Whitney test, unpaired, two-tailed). This was consistent with the previous observation from *reeler* mice (Stanfield and Cowan, 1979b), and indicated that there was no catastrophic loss of the proliferating cell population at this stage of dentate gyrus development. The fraction of cells that remained in cell cycle (BrdU+,Ki67+/Total BrdU) was reduced (Figure 2C, p=0.0022, Mann-Whitney test, unpaired, two-tailed), suggesting a mild neurogenesis defect. In addition, there were very few cells labeled with anti-activated caspase-3 in the entire hippocampus at this age (Supporting figure 1), suggesting that there is no significant increase in cell death.

### Distribution of Tbr2-positive intermediate progenitor cells is abnormal

To further understand abnormalities in the dentate gyrus development, we examined the distribution of Tbr2-positive cells (Figure 3). Tbr2 marks the intermediate neuronal progenitors, which generate the majority of granule neurons in the dentate gyrus (Hodge et al., 2012). We observed an abnormal distribution of Tbr2-positive cells at all ages examined.

**Figure 3.**
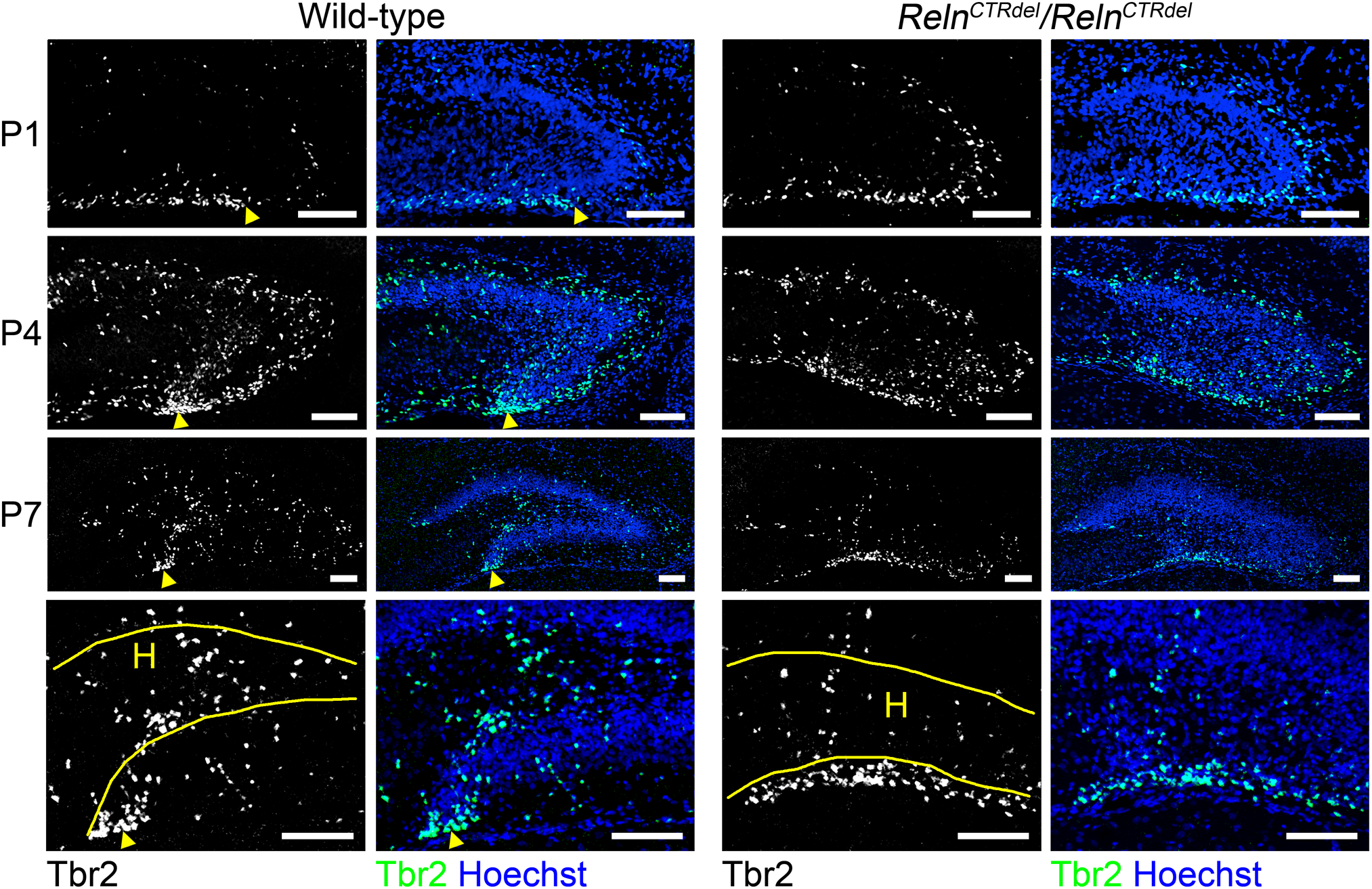
Lack of intermediate progenitor accumulation at the fimbriodentate junction. In wild-type mice, Tbr2-positive cells (gray or green) accumulate at the fimbriodentate junction (arrowheads) starting at P1 (n=2 each) and more clearly at P4 (n=2 each) and P7 (n=2 each). This accumulation at FDJ is less apparent in the mutant, and the labeled cells are spread out along the ventricular surface. Enlarged imaged of the hilus at P7 are shown in the lower panel. *Reln*^*CTRdel*^ mice have fewer Tbr2-labeled cells in the hilus (H) compared to wild-type mice; it appears as if many cells are still confined to the ventricular surface. Nuclei were stained using Hoechst (blue). Scale bars, 100 μm.

The wild-type mice start to show an accumulation of Tbr2-positive cells near the fimbriodentate junction (Figure 3, arrowheads) at P1; however, this was not apparent in the mutant. The cluster of Tbr2-positive cells at the fimbriodentate junction became more evident at P4, and was still rudimentary in the mutant. This lack of the intermediate progenitor accumulation resembled the lack of the BrdU or Ki67-positive neurogenic cluster shown in Figure 2, and it is likely that the Tbr2-positive cells that accumulate at the fimbriodentate junction comprise the neurogenic cluster together. In wild-type mice at P7, the cluster of Tbr2-positive cells was still present at the fimbriodentate junction (Figure 3). In the mutant, Tbr2-positive cells along the subpial surface were more widely scattered than in wild-type, although a rudimentary fimbriodentate junction seemed visible. It appeared that fewer Tbr2-positive cells were present in the hilus of the mutant than that of wild-type mice at P7, suggesting a change in the migration of progenitor cells.

### Granule neuron precursor distribution in the hilus is abnormal

We also examined Prox1 or NeuroD-positive populations to determine whether the mutant has a granule neuron production defect (Figure 4). At P7, the number of cells expressing these markers of more differentiated progenitors was greater in the hilus of the mutant. These results suggest that the progenitor population in the hilus is altered in the mutant. In addition, there are more Prox1 or NeuroD-positive cells in the mutant SPB, which appears thicker than the wild-type SPB.

**Figure 4.**
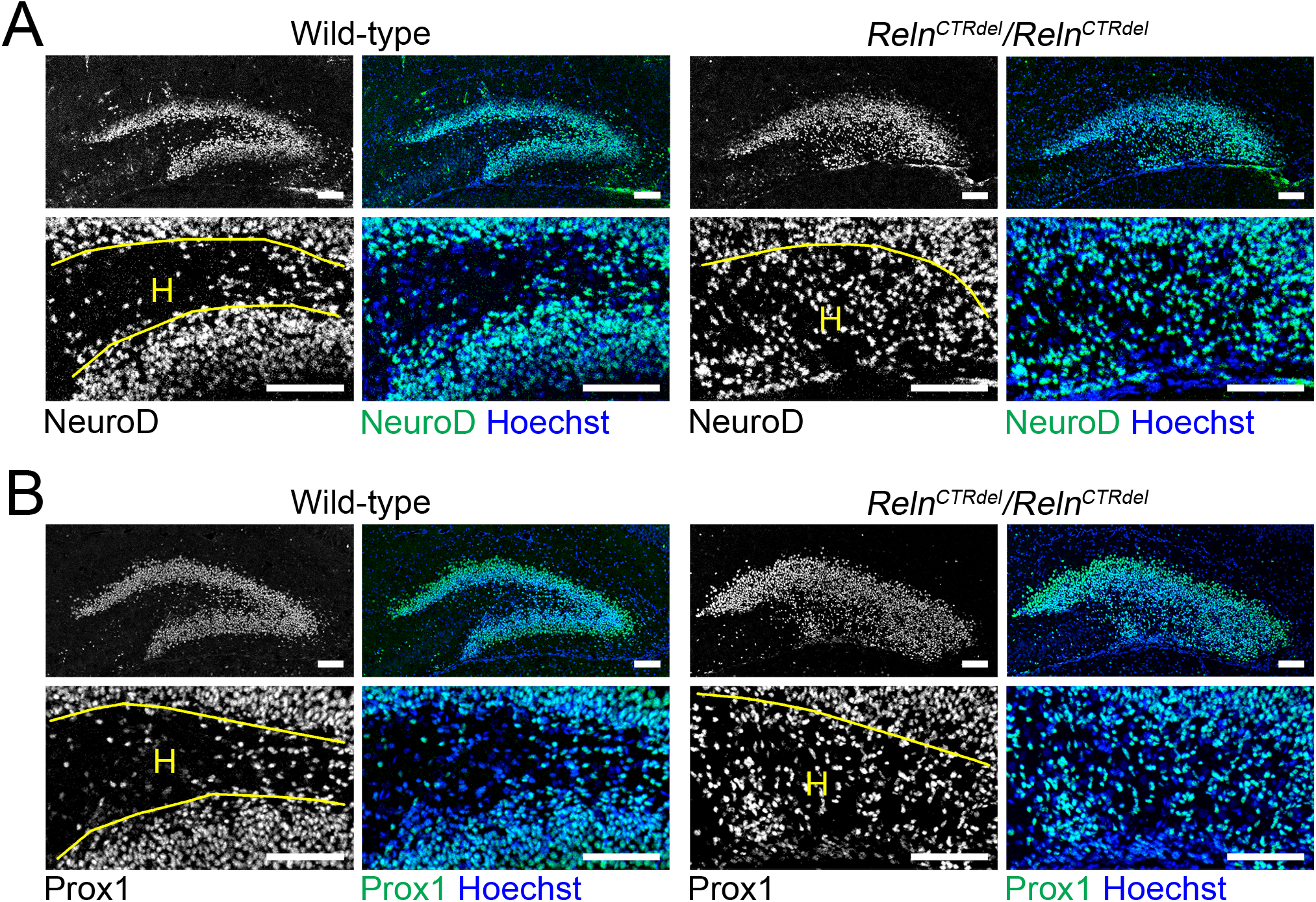
Abnormal distribution of the granule neurons. Granule neuron precursor markers (gray or green), NeuroD (**A**) and Prox1 (**B**), staining indicate the presence of abnormal progenitor population in the hilus (H) of *Reln*^*CTRdel*^ at P7 (n=2 each). Nuclei were stained using Hoechst (blue). Enlarged images are shown in lower panels. Scale bars, 100 μm.

### Radial glial scaffold defect is specific to the IPB

In rodents, the primary radial glial scaffold undergoes extensive reorganization soon after birth, and is augmented by the addition of new (secondary) radial fibers in the dentate gyrus (Rickmann et al., 1987; Brunne et al., 2010). Mice lacking RELN or DAB1 have extreme disorganization of the secondary radial glial scaffold, resulting in dispersion of granule cells in the hilus (Forster et al., 2002). Therefore, we examined the radial glial scaffold in the dentate gyrus of *Reln*^*CTRdel*^ mutants using anti-GFAP and anti-BLBP antibodies (Figure 5).

**Figure 5.**
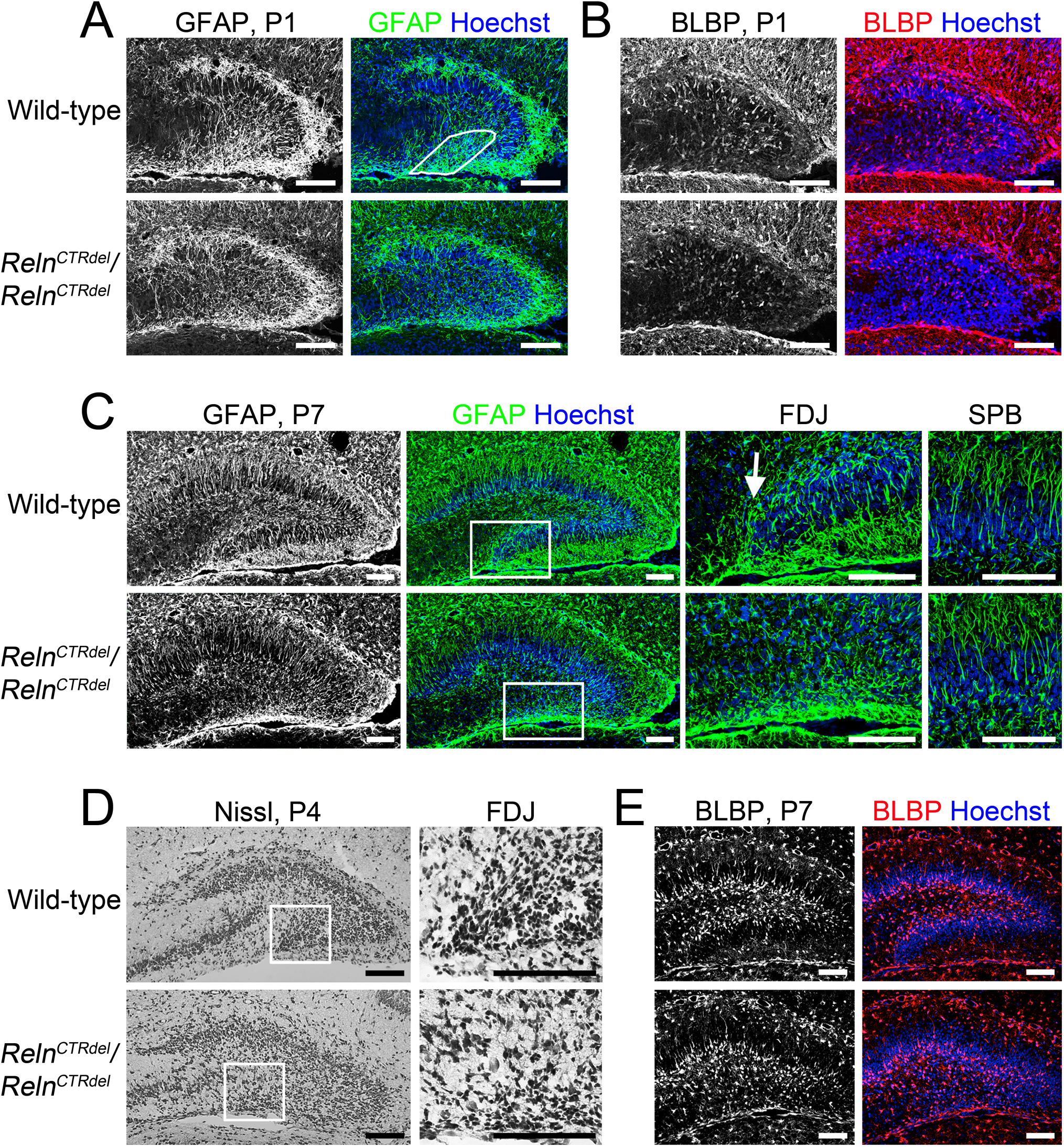
Lack of the secondary glial scaffold formation in the IPB. **A:** At P1 when the secondary glial scaffold formation begins in the IPB, the mutant mice lack dense GFAP-stained fibers (gray or green, marked region) that are apparent near the fimbriodentate junction of wild-type mice (n=2 each). Note the premature secondary glial scaffold can be already seen in the SPB of both wild-type and mutant. **B:** At P1, BLBP-positive cell bodies (gray or red) appear near the IPB in wild-type mice (n=2 each). Fewer BLBP-positive cells are found near the IPB of *Reln*^*CTRdel*^ mice. **C:** Images of GFAP-stained (gray or green) dentate gyrus at P7 (left, n=2 each). Enlarged images of the boxed fimbriodentate junction (FDJ) regions are shown in the center. In wild-type mice, GFAP-positive fibers are surrounding the growing tip of the infrapyramidal blade at the fimbriodentate junction (arrow). This structure is absent in the mutant. The secondary glia scaffold in the SPB is relatively normal (right, higher-magnification images). **D:** At P4, a stream of migrating cells with elongated morphology is absent in the mutant fimbriodentate junction (n=1 each). Enlarged images of the fimbriodentate junction regions are shown on the right. Nissl stain. **E:** At P7, abnormal distribution of BLBP-positive cells (gray or red) in the hilus become more evident (n=2 each). The cells are directed toward the SPB in the mutant. Nuclei were stained using Hoechst (blue). Scale bars, 100 μm. Enlarged images of A (color panel), B (color panel), D (right panel), and E (color panel) can be found in Supporting figure 2.

We observed abnormal development of the secondary glial scaffold in *Reln*^*CTRdel*^ mutants. The secondary glial scaffold begins to be radially organized in the SPB by P1 (Figure 5A), as previously described (Forster et al., 2002; Brunne et al., 2013). Similarly, some BLBP-positive cells were already a part of the SPB secondary glial scaffold (Figure 5B). In the wild-type IPB, short densely organized GFAP-stained fibers were seen near the fimbriodentate junction (Figure 5A, marked region). In *Reln*^*CTRdel*^, however, this pattern of glia fibers was not seen at the fimbriodentate junction.

By P7, the mutant fully developed the secondary glial scaffold in the SPB comparable with the wild-type (Figure 5C, left and right), but this was completely absent in the IPB (Figure 5C, left). In the IPB, densely organized glial fibers at the fimbriodentate junction remained absent (Figure 5C, center). These observations suggest that normal RELN function is required for glial fiber organization at the fimbriodentate junction, and CTR truncation disrupts this function.

As seen on Nissl stain, the cells at the fimbriodentate junction in the wild-type had elongated shape, suggestive of glial-guided migration from the fimbriodentate junction to the hilus (Figure 5D). However, the fimbriodentate junction in *Reln*^*CTRdel*^ did not contain cells with elongated shape. This observation is consistent with our speculation that the progenitor cells are confined to the infrapyramidal subpial surface in the mutant.

The abnormal distribution of BLBP-positive cells was obvious at P7. BLBP-positive cells were found in the subgranular zone in the wild-type dentate gyrus, but the infrapyramidal subgranular zone was absent in *Reln*^*CTRdel*^ mutant due to truncation of the IPB (Figure 5E). This phenotype, in addition to abnormal distribution of Tbr2, NeuroD, or Prox1-positive cells, may indicate that organization of the hilar and subgranular neurogenic zones is disturbed in the mutant mice.

Altogether, these results suggest that local abnormalities in the secondary radial glial scaffold may lead to abnormal progenitor cell migration at the fimbriodentate junction and neurogenic zone organization, ultimately resulting in an absent IPB.

## DISCUSSION

### A *Reln* mutation can cause abnormal positioning of Cajal-Retzius cells

Cajal-Retzius cells and reelin signaling play an important role in development of hippocampal laminar organization and connections (Del Rio et al., 1997; Super et al., 1998; Borrell et al., 1999a; Borrell et al., 1999b; Borrell et al., 2007). In *reeler* mutant mice, the dentate gyrus retains a laminated outer molecular layer, in which entorhinal afferents terminate (Deller et al., 1999). These mice appear to form both the SPB and IPB, although they have a highly disorganized granule cell layer. Calretinin-positive Cajal-Retzius cells are present in both SPB and IPB of *reeler* mice (Deller et al., 1999). Cajal-Retzius cells are known to be the transient target for the entorhinal projection (Del Rio et al., 1997), accounting for the presence of the molecular layer in *reeler* mice.

*Reln*^*CTRdel*^ mice do not form the granule cell layer nor the molecular layer in the IPB (Ha et al., 2017). We observed that fewer Cajal-Retzius cells were present in the subpial surface of the prospective IPB (Figure 1). The lack of the target cells in IPB may lead to a failure of entorhinal projection innervation, and this would account for the absence of the infrapyramidal molecular layer in *Reln*^*CTRdel*^ mice. On the contrary, the SPB has Cajal-Retzius cells and a molecular layer, suggesting that the entorhinal afferents innervation does occur.

Our phenotypic analysis provides direct genetic evidence that a *Reln* mutation can cause abnormal positioning of Cajal-Retzius cells, which has not previously been shown for any other allele of *Reln*. Different subtypes of Cajal-Retzius cells arise from multiple sources and migrate tangentially (Reviewed in Barber and Pierani, 2016). The cortical hem, immediate adjacent and medial to the dentate gyrus, is one of the main sources of Cajal-Retzius cells (Grove et al., 1998; Meyer et al., 2002; Takiguchi-Hayashi et al., 2004; Yoshida et al., 2006). The majority of hippocampal Cajal-Retzius cells originate from the cortical hem, and they migrate along the fimbrial BLBP-positive glial fibers to reach the dentate gyrus (Gu et al., 2011). Increased Cajal-Retzius cell number in the SPB of *Reln*^*CTRdel*^ mice suggests a possibility that Cajal-Retzius cells might overmigrate to the SPB, instead of stopping at their normally destined position within the subpial surface of IPB.

The mechanism by which CTR-truncated RELN mediates abnormal positioning of the Cajal-Retzius cells is not obvious. Potential mechanisms involved could be altered CXCR4-CXCL12 signaling (Bagri et al., 2002; Lu et al., 2002; Li et al., 2009; Hodge et al., 2013; Parisot et al., 2017), and an abnormal contact-mediated repulsion, which was shown to be mediated by Eph/ephrin signaling (Villar-Cervino et al., 2013).

### Defects in the secondary glial scaffold and neurogenic niche at the fimbriodentate junction result in truncation of the infrapyramidal blade

*Reeler* does not develop the secondary glial scaffold at all, and a lack of this structure has been considered as a main cause of dispersed granule cell layer in the dentate gyrus (Forster et al., 2002; Frotscher et al., 2003; Weiss et al., 2003). In addition, it was shown that some Tbr2-positive cells in the suprapyramidal subpial zone of the SPB migrate inward, crossing the granule cell layer (Li et al., 2009). The secondary glial fibers in the SPB provide the substrate for this migration. *Reeler* mice have a defect in this process, and the Tbr2-positive neurons are retained in the suprapyramidal subpial zone (Li et al., 2009). *Reln*^*CTRdel*^ homozygotes have the secondary glial scaffold in the SPB (Figure 5) and did not show abnormal subpial-subgranular transition (Figure 3).

However, *Reln*^*CTRdel*^ does not form the secondary glial scaffold in the IPB (Figure 5), where the RELN-expressing cells are absent. Abnormal organization of glial fibers at the fimbriodentate junction observed soon after birth is the early structural defect preceding the formation of infrapyramidal granule cell layer (Figures 5A, B). This suggests that a defect in the radial glial scaffold, which normally provides a migratory frame for progenitor cells, might be a cause of the dentate gyrus malformation in *Reln*^*CTRdel*^. A similar defect was previously reported in *Foxg1* mutant mice lacking the IPB (Tian et al., 2012), suggesting the importance of the glial fibers at the fimbriodentate junction for the formation of the IPB. It has been shown that RELN mediates secondary glial scaffold development by amplifying Notch signaling (Sibbe et al., 2009).

We observed no difference in total BrdU-labeled cell number in the entire area of the dentate gyrus, with only a minor neurogenesis defect (Figure 2). In other mutant mice that have a severe neurogenesis defect, both blades are reduced in length and thickness (Lu et al., 2002; Li et al., 2009; Tian et al., 2012; Hodge et al., 2013). As *Reln*^*CTRdel*^ have a relatively well-developed SPB, it is possible that truncation of the IPB originates from a local defect of radial glia-like neural stem cells, as well as granule neurons. The abnormal distribution of dividing cells (Figure 2) and intermediate progenitors (Figure 3) in the mutant suggests that a neurogenic niche at the fimbriodentate junction is critical for the extension of IPB. Without proper organization at the fimbriodentate junction, progenitors and dividing cells fail to contribute to the IPB formation.

Based on the above observations, we propose a model to explain disorganization of the neurogenic niche at the fimbriodentate junction (Figure 6, left panel), and the potential role of RELN in this process. During the formation of the normal IPB, a subpopulation of progenitor cells accumulates at the fimbriodentate junction (Figures 2 and 3). From the fimbriodentate junction, these progenitors are guided to the inner surface of the IPB by short, densely organized, curved glial fibers (Figures 5). They may divide again in close proximity at the tip of growing IPB, and these cells altogether would appear as a neurogenic cluster. Concurrently, Cajal-Retzius cells migrate into the hippocampal fissure first, and then later-arriving cells cover the subpial surface of the IPB. Localization of Cajal-Retzius cells in the IPB correlates with the IPB formation spatially and temporally, and may be related to the addition of new radial units (radial glia-like scaffold cells plus granule neurons) to the IPB. Our observations suggest that proper localization of Cajal-Retzius cells is required to establish glial fiber organization and neurogenic niche at the fimbriodentate junction (Figure 5).

**Figure 6.**
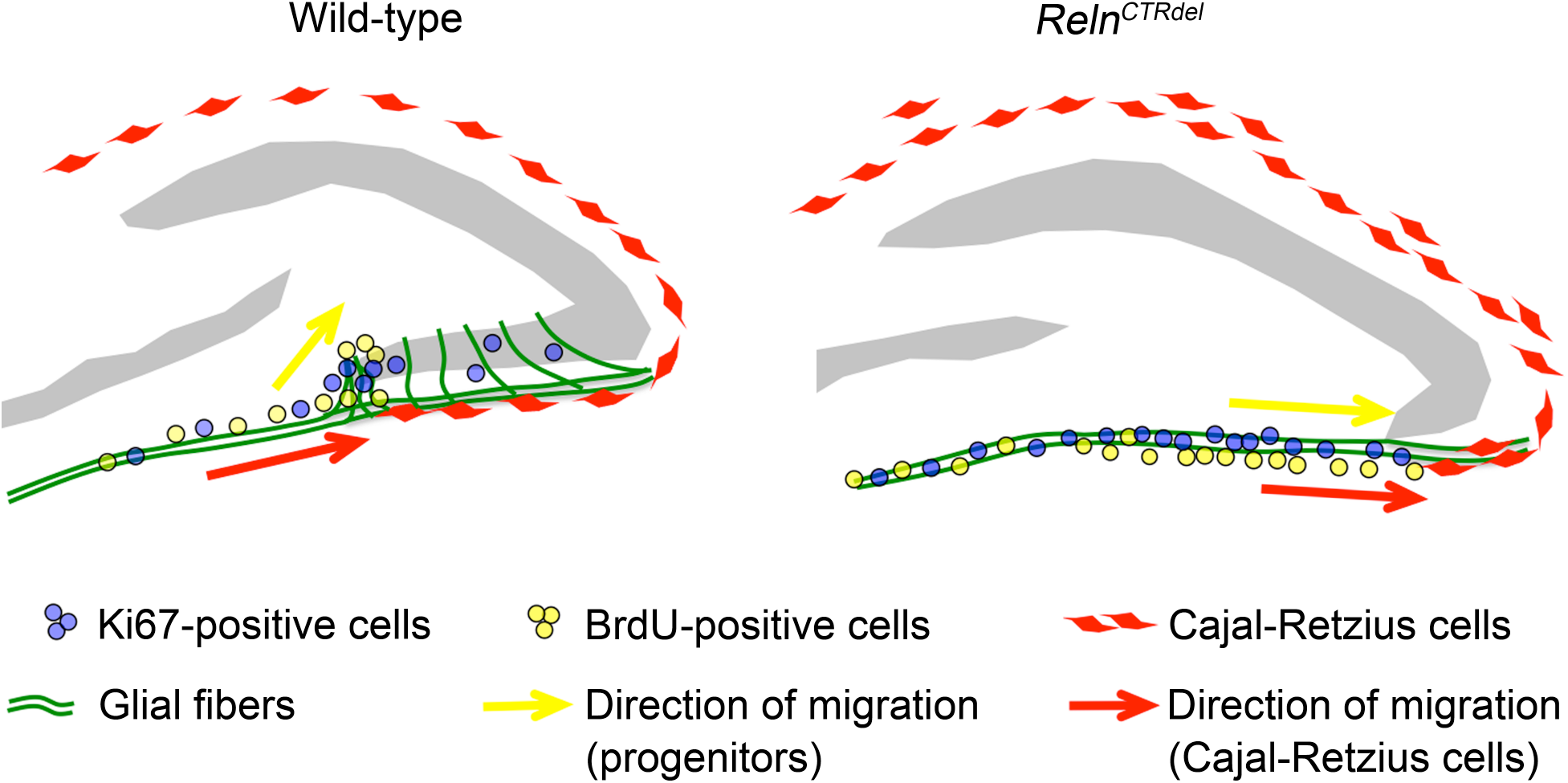
A model of the infrapyramidal malformation. During the first postnatal week, Cajal-Retzius cells disperse along the subpial surface of the IPB. The progenitors migrate along the dentate migratory stream, then along the densely organized glia scaffold at the fimbriodentate junction, proliferate at the tip and extend the IPB. In *Reln*^*CTRdel*^ mutant, Cajal-Retzius cells overmigrate to the SPB and are absent in the subpial surface of the IPB. Without proper RELN signal, the glia scaffolds cannot become organized at the fimbriodentate junction, the progenitor cells fail to migrate toward the hilus, and proliferate within the subpial surface. The mutant dentate gyrus fails to form the outer shell in the IPB, which affects subsequent neurogenesis events in the dentate gyrus.

In *Reln*^*CTRdel*^ mutant, Cajal-Retzius cells fail to settle in the IPB subpial surface, the glial scaffold at the fimbriodentate junction is not organized, this prevent transhilar migration of progenitors, and the mutants fail to organize a neurogenic cluster (Figure 6, right panel). The outer shell of the dentate gyrus granule cell layer is shaped during the first postnatal week (Altman and Bayer, 1990a); once this critical period is missed granule neurons born later from the hilus and subgranular neurogenic zone cannot compensate for the loss of the IPB, and further growth of the IPB does not occur.

### Does reelin act as a repulsive signal at the fimbriodentate junction?

*Reln* act as both an attractive (or permissive) signal and a stop signal for cortical neurons (Ogawa et al., 1995; Frotscher, 1997; Schiffmann et al., 1997; Sheppard and Pearlman, 1997; Dulabon et al., 2000; Herrick and Cooper, 2002; Olson et al., 2006; Hack et al., 2007; Zhao and Frotscher, 2010). It is thought that APOER2 is important for the former, and VLDLR is required for the latter. However, the granule neurons can reach the dentate gyrus without an attractive signal in *reeler* mice, and the role of RELN as a positional cue for hippocampal development has been more focused on the glia scaffold organization and formation of the compact granule cell layers (Frotscher et al., 2003; Forster et al., 2006; Zhao and Frotscher, 2010). In addition to the above model, we further propose that RELN might function as a repulsive signal locally at the fimbriodentate junction. CTR-truncation, if it disrupts the repulsive function, might cause Cajal-Retzius cell overmigration toward the SPB and disable accumulation of granule neuron progenitors and their transhilar migration at the fimbriodentate junction (Figure 6, right panel).

### Can Differential Receptor Binding Account For The IPB Defect?

We have demonstrated that a CTR-truncation of RELN disrupts an interaction with VLDLR (Ha et al., 2017). It is challenging to interpret the IPB defect in the context of differential RELN receptor binding, since *Vldlr* null mice do not show an obvious dentate gyrus malformation. Interestingly, double-homozygote mice carrying both *Reln*^*CTRdel*^ and *Vldlr* null mutations display a more severe IPB defect than *Reln*^*CTRdel*^ mice (Ha et al., 2017), possibly because abnormal positioning of the Cajal-Retzius cells and neuronal progenitors is exacerbated. This additive effect suggests the possibility that disrupted binding to VLDLR is involved in the IPB defect, and also that VLDLR may not be the only receptor involved. We speculate that CTR-truncated RELN interacts abnormally with other RELN receptor(s), which might be mediating the repulsive signal at the fimbriodentate junction. In double-homozygote mice, the complete absence of VLDLR interaction would further increase unbound CTR-truncated RELN protein, which would exacerbate aberrant interactions with other RELN receptor(s). This hypothesis would explain why *Vldlr* null mice have a normal IPB, as they express normal RELN proteins, which would not cause aberrant interactions. This would also account for the absence of a specific IPB defect in RELN-deficient mice; they do not express RELN, so aberrant interactions cannot occur.

One can further speculate regarding the nature of these hypothetical aberrant interactions. Firstly, we have previously discussed a possibility that a subtle change in the interaction of CTR-truncated RELN and APOER2 might contribute to the hippocampal abnormality (Ha et al., 2017). *Apoer2* null mice display morphological defects in the dentate gyrus that include an abnormal IPB (Fish and Krucker, 2008; also see Supporting figure 3), which has not been previously emphasized. Of note, the double homozygote of *Reln*^*CTRdel*^ and *Apoer2* null mutations appear identical to *reeler* with an intact outer molecular layer in the IPB (Ha et al., 2017), suggesting the IPB defect is mitigated by the absence of APOER. Alternatively, it has been suggested that there exists an unrecognized APOER2/VLDLR-associated coreceptor that can bind to the CTR domain of RELN and is required to fully activate reelin signaling (Nakano et al., 2007). Also, mice lacking beta1-integrin, which can bind to RELN (Dulabon et al., 2000), display a local secondary glial scaffold disruption at the fimbriodentate junction, although they still develop an IPB (Forster et al., 2002). Perhaps aberrant binding to more than a single RELN receptor mediates the hippocampal defects we see in *Reln*^*CTRdel*^ mice.

### Potential clinical relevance

In addition to type II lissencephaly with cerebellar hypoplasia, *RELN* mutations are also associated with autosomal-dominant lateral temporal-lobe epilepsy (ADLTE) with reduced penetrance (Dazzo et al., 2015). Morphological abnormalities were not identifiable from conventional MRI scans of ADLTE patients carrying *RELN* mutations (Michelucci et al., 2017), and the pathophysiology of the disorder remains uncertain. The observation that *RELN* mutations can be associated with hypomorphic phenotypes in human patients suggests that characterization of hypomorphic *Reln* mutant mice may benefit our understanding of human disorders.

## OTHER ACKNOWLEDGEMENTS

We thank Anna M. Lindsay and Dr. Scott Houghtaling for technical assistance.

## REFERENCES

Altman J, Bayer SA. 1990a. Migration and distribution of two populations of hippocampal granule cell precursors during the perinatal and postnatal periods. J Comp Neurol 301:365–381. doi:10.1002/cne.903010304

Altman J, Bayer SA. 1990b. Mosaic organization of the hippocampal neuroepithelium and the multiple germinal sources of dentate granule cells. J Comp Neurol 301:325–342. doi:10.1002/cne.903010302

Bagri A, Gurney T, He X, Zou YR, Littman DR, Tessier-Lavigne M, Pleasure SJ. 2002. The chemokine SDF1 regulates migration of dentate granule cells. Development 129:4249–4260.

Barber M, Pierani A. 2016. Tangential migration of glutamatergic neurons and cortical patterning during development: Lessons from Cajal-Retzius cells. Dev Neurobiol 76:847–881. doi:10.1002/dneu.22363

Borrell V, Del Rio JA, Alcantara S, Derer M, Martinez A, D’Arcangelo G, Nakajima K, Mikoshiba K, Derer P, Curran T, Soriano E. 1999a. Reelin regulates the development and synaptogenesis of the layer-specific entorhino-hippocampal connections. J Neurosci 19:1345–1358.

Borrell V, Ruiz M, Del Rio JA, Soriano E. 1999b. Development of commissural connections in the hippocampus of reeler mice: evidence of an inhibitory influence of Cajal-Retzius cells. Exp Neurol 156:268–282.

Borrell V, Pujadas L, Simo S, Dura D, Sole M, Cooper JA, Del Rio JA, Soriano E. 2007. Reelin and mDab1 regulate the development of hippocampal connections. Mol Cell Neurosci 36:158–173. doi:10.1016/j.mcn.2007.06.006

Brunne B, Zhao S, Derouiche A, Herz J, May P, Frotscher M, Bock HH. 2010. Origin, maturation, and astroglial transformation of secondary radial glial cells in the developing dentate gyrus. Glia 58:1553–1569. doi:10.1002/glia.21029

Brunne B, Franco S, Bouche E, Herz J, Howell BW, Pahle J, Muller U, May P, Frotscher M, Bock HH. 2013. Role of the postnatal radial glial scaffold for the development of the dentate gyrus as revealed by Reelin signaling mutant mice. Glia 61:1347–1363. doi:10.1002/glia.22519

D’Arcangelo G, Nakajima K, Miyata T, Ogawa M, Mikoshiba K, Curran T. 1997. Reelin is a secreted glycoprotein recognized by the CR-50 monoclonal antibody. J Neurosci 17:23–31.

Dazzo E, Fanciulli M, Serioli E, Minervini G, Pulitano P, Binelli S, Di Bonaventura C, Luisi C, Pasini E, Striano S, Striano P, Coppola G, Chiavegato A, Radovic S, Spadotto A, Uzzau S, La Neve A, Giallonardo AT, Mecarelli O, Tosatto SC, Ottman R, Michelucci R, Nobile C. 2015. Heterozygous reelin mutations cause autosomal-dominant lateral temporal epilepsy. Am J Hum Genet 96:992–1000. doi:10.1016/j.ajhg.2015.04.020

de Bergeyck V, Nakajima K, Lambert de Rouvroit C, Naerhuyzen B, Goffinet AM, Miyata T, Ogawa M, Mikoshiba K. 1997. A truncated Reelin protein is produced but not secreted in the ‘Orleans’ reeler mutation (*Reln ^rl-Orl^*). Brain Res Mol Brain Res 50:85–90. doi:10.1016/S0169-328X(97)00166-6

de Bergeyck V, Naerhuyzen B, Goffinet AM, Lambert de Rouvroit C. 1998. A panel of monoclonal antibodies against reelin, the extracellular matrix protein defective in reeler mutant mice. J Neurosci Methods 82:17–24.

Del Rio JA, Heimrich B, Borrell V, Forster E, Drakew A, Alcantara S, Nakajima K, Miyata T, Ogawa M, Mikoshiba K, Derer P, Frotscher M, Soriano E. 1997. A role for Cajal-Retzius cells and reelin in the development of hippocampal connections. Nature 385:70–74. doi:10.1038/385070a0

Deller T, Drakew A, Frotscher M. 1999. Different primary target cells are important for fiber lamination in the fascia dentata: a lesson from reeler mutant mice. Exp Neurol 156:239–253. doi:10.1006/exnr.1999.7020

Dulabon L, Olson EC, Taglienti MG, Eisenhuth S, McGrath B, Walsh CA, Kreidberg JA, Anton ES. 2000. Reelin binds alpha3beta1 integrin and inhibits neuronal migration. Neuron 27:33–44. doi:10.1016/S0896-6273(00)00007-6

Fish KN, Krucker T. 2008. Functional consequences of hippocampal neuronal ectopia in the apolipoprotein E receptor-2 knockout mouse. Neurobiol Dis 32:391–401. doi:10.1016/j.nbd.2008.07.023

Forster E, Tielsch A, Saum B, Weiss KH, Johanssen C, Graus-Porta D, Muller U, Frotscher M. 2002. Reelin, Disabled 1, and beta 1 integrins are required for the formation of the radial glial scaffold in the hippocampus. Proc Natl Acad Sci U S A 99:13178–13183. doi:10.1073/pnas.202035899

Forster E, Zhao S, Frotscher M. 2006. Laminating the hippocampus. Nat Rev Neurosci 7:259–267. doi:10.1038/nrn1882

Frotscher M. 1997. Dual role of Cajal-Retzius cells and reelin in cortical development. Cell Tissue Res 290:315–322. doi:10.1007/s004410050936

Frotscher M, Haas CA, Forster E. 2003. Reelin controls granule cell migration in the dentate gyrus by acting on the radial glial scaffold. Cereb Cortex 13:634–640.

Grove EA, Tole S, Limon J, Yip L, Ragsdale CW. 1998. The hem of the embryonic cerebral cortex is defined by the expression of multiple Wnt genes and is compromised in Gli3-deficient mice. Development 125:2315–2325.

Gu X, Liu B, Wu X, Yan Y, Zhang Y, Wei Y, Pleasure SJ, Zhao C. 2011. Inducible genetic lineage tracing of cortical hem derived Cajal-Retzius cells reveals novel properties. PLoS One 6:e28653. doi:10.1371/journal.pone.0028653

Ha S, Stottmann RW, Furley AJ, Beier DR. 2015. A forward genetic screen in mice identifies mutants with abnormal cortical patterning. Cereb Cortex 25:167–179. doi:10.1093/cercor/bht209

Ha S, Tripathi PP, Mihalas AB, Hevner RF, Beier DR. 2017. C-Terminal Region Truncation of RELN Disrupts an Interaction with VLDLR, Causing Abnormal Development of the Cerebral Cortex and Hippocampus. J Neurosci 37:960–971. doi:10.1523/JNEUROSCI.1826-16.2017

Hack I, Hellwig S, Junghans D, Brunne B, Bock HH, Zhao S, Frotscher M. 2007. Divergent roles of ApoER2 and Vldlr in the migration of cortical neurons. Development 134:3883–3891. doi:10.1242/dev.005447

Hartmann D, Sievers J, Pehlemann FW, Berry M. 1992. Destruction of meningeal cells over the medial cerebral hemisphere of newborn hamsters prevents the formation of the infrapyramidal blade of the dentate gyrus. J Comp Neurol 320:33–61. doi:10.1002/cne.903200103

Hartmann D, Frotscher M, Sievers J. 1994. Development of granule cells, and afferent and efferent connections of the dentate gyrus after experimentally induced reorganization of the supra- and infrapyramidal blades. Acta Anat (Basel) 150:25– 37.

Hayashi K, Kubo K, Kitazawa A, Nakajima K. 2015. Cellular dynamics of neuronal migration in the hippocampus. Front Neurosci 9:135. doi:10.3389/fnins.2015.00135

Herrick TM, Cooper JA. 2002. A hypomorphic allele of dab1 reveals regional differences in reelin-Dab1 signaling during brain development. Development 129:787–796.

Hodge RD, Kowalczyk TD, Wolf SA, Encinas JM, Rippey C, Enikolopov G, Kempermann G, Hevner RF. 2008. Intermediate progenitors in adult hippocampal neurogenesis: Tbr2 expression and coordinate regulation of neuronal output. J Neurosci 28:3707–3717. doi:10.1523/JNEUROSCI.4280-07.2008

Hodge RD, Nelson BR, Kahoud RJ, Yang R, Mussar KE, Reiner SL, Hevner RF. 2012. Tbr2 is essential for hippocampal lineage progression from neural stem cells to intermediate progenitors and neurons. J Neurosci 32:6275–6287. doi:10.1523/JNEUROSCI.0532-12.2012

Hodge RD, Garcia AJ, 3rd, Elsen GE, Nelson BR, Mussar KE, Reiner SL, Ramirez JM, Hevner RF. 2013. Tbr2 expression in Cajal-Retzius cells and intermediate neuronal progenitors is required for morphogenesis of the dentate gyrus. J Neurosci 33:4165–4180. doi:10.1523/JNEUROSCI.4185-12.2013

Kohno T, Honda T, Kubo K, Nakano Y, Tsuchiya A, Murakami T, Banno H, Nakajima K, Hattori M. 2015. Importance of reelin C-terminal region in the development and maintenance of the postnatal cerebral cortex and its regulation by specific proteolysis. J Neurosci 35:4776–4787. doi:10.1523/JNEUROSCI.4119-14.2015

Li G, Pleasure SJ. 2005. Morphogenesis of the dentate gyrus: what we are learning from mouse mutants. Dev Neurosci 27:93–99. doi:10.1159/000085980

Li G, Kataoka H, Coughlin SR, Pleasure SJ. 2009. Identification of a transient subpial neurogenic zone in the developing dentate gyrus and its regulation by Cxcl12 and reelin signaling. Development 136:327–335. doi:10.1242/dev.025742

Lu M, Grove EA, Miller RJ. 2002. Abnormal development of the hippocampal dentate gyrus in mice lacking the CXCR4 chemokine receptor. Proc Natl Acad Sci U S A 99:7090–7095. doi:10.1073/pnas.092013799

Meyer G, Perez-Garcia CG, Abraham H, Caput D. 2002. Expression of p73 and Reelin in the developing human cortex. J Neurosci 22:4973–4986.

Michelucci R, Pulitano P, Di Bonaventura C, Binelli S, Luisi C, Pasini E, Striano S, Striano P, Coppola G, La Neve A, Giallonardo AT, Mecarelli O, Serioli E, Dazzo E, Fanciulli M, Nobile C. 2017. The clinical phenotype of autosomal dominant lateral temporal lobe epilepsy related to reelin mutations. Epilepsy Behav 68:103–107. doi:10.1016/j.yebeh.2016.12.003

Nakamura K, Beppu M, Sakai K, Yagyu H, Matsumaru S, Kohno T, Hattori M. 2016. The C-terminal region of Reelin is necessary for proper positioning of a subset of Purkinje cells in the postnatal cerebellum. Neuroscience 336:20–29. doi:10.1016/j.neuroscience.2016.08.039

Nakano Y, Kohno T, Hibi T, Kohno S, Baba A, Mikoshiba K, Nakajima K, Hattori M. 2007. The extremely conserved C-terminal region of Reelin is not necessary for secretion but is required for efficient activation of downstream signaling. J Biol Chem 282:20544–20552. doi:10.1074/jbc.M702300200

Nicola Z, Fabel K, Kempermann G. 2015. Development of the adult neurogenic niche in the hippocampus of mice. Front Neuroanat 9:53. doi:10.3389/fnana.2015.00053

Ogawa M, Miyata T, Nakajima K, Yagyu K, Seike M, Ikenaka K, Yamamoto H, Mikoshiba K. 1995. The reeler gene-associated antigen on Cajal-Retzius neurons is a crucial molecule for laminar organization of cortical neurons. Neuron 14:899–912. doi:10.1016/0896-6273(95)90329-1

Olson EC, Kim S, Walsh CA. 2006. Impaired neuronal positioning and dendritogenesis in the neocortex after cell-autonomous Dab1 suppression. J Neurosci 26:1767–1775. doi:10.1523/JNEUROSCI.3000-05.2006

Parisot J, Flore G, Bertacchi M, Studer M. 2017. COUP-TFI mitotically regulates production and migration of dentate granule cells and modulates hippocampal Cxcr4 expression. Development 144:2045–2058. doi:10.1242/dev.139949

Pedroni A, Minh do D, Mallamaci A, Cherubini E. 2014. Electrophysiological characterization of granule cells in the dentate gyrus immediately after birth. Front Cell Neurosci 8:44. doi:10.3389/fncel.2014.00044

Pesold C, Impagnatiello F, Pisu MG, Uzunov DP, Costa E, Guidotti A, Caruncho HJ. 1998. Reelin is preferentially expressed in neurons synthesizing gammaaminobutyric acid in cortex and hippocampus of adult rats. Proc Natl Acad Sci U S A 95:3221–3226.

Rickmann M, Amaral DG, Cowan WM. 1987. Organization of radial glial cells during the development of the rat dentate gyrus. J Comp Neurol 264:449–479. doi:10.1002/cne.902640403

Sakai K, Shoji H, Kohno T, Miyakawa T, Hattori M. 2016. Mice that lack the C-terminal region of Reelin exhibit behavioral abnormalities related to neuropsychiatric disorders. Sci Rep 6:28636. doi:10.1038/srep28636

Schiffmann SN, Bernier B, Goffinet AM. 1997. Reelin mRNA expression during mouse brain development. Eur J Neurosci 9:1055–1071. doi:10.1111/j.1460-9568.1997.tb01456.x

Sheppard AM, Pearlman AL. 1997. Abnormal reorganization of preplate neurons and their associated extracellular matrix: an early manifestation of altered neocortical development in the reeler mutant mouse. J Comp Neurol 378:173–179. doi:10.1002/(SICI)1096-9861(19970210)378:2<173::AID-CNE2>3.0.CO;2-0

Sibbe M, Forster E, Basak O, Taylor V, Frotscher M. 2009. Reelin and Notch1 cooperate in the development of the dentate gyrus. J Neurosci 29:8578–8585. doi:10.1523/JNEUROSCI.0958-09.2009

Stanfield BB, Cowan WM. 1979a. The morphology of the hippocampus and dentate gyrus in normal and reeler mice. J Comp Neurol 185:393–422. doi:10.1002/cne.901850302

Stanfield BB, Cowan WM. 1979b. The development of the hippocampus and dentate gyrus in normal and reeler mice. J Comp Neurol 185:423–459. doi:10.1002/cne.901850303

Super H, Martinez A, Del Rio JA, Soriano E. 1998. Involvement of distinct pioneer neurons in the formation of layer-specific connections in the hippocampus. J Neurosci 18:4616–4626.

Takiguchi-Hayashi K, Sekiguchi M, Ashigaki S, Takamatsu M, Hasegawa H, Suzuki-Migishima R, Yokoyama M, Nakanishi S, Tanabe Y. 2004. Generation of reelin-positive marginal zone cells from the caudomedial wall of telencephalic vesicles. J Neurosci 24:2286–2295. doi:10.1523/JNEUROSCI.4671-03.2004

Tian C, Gong Y, Yang Y, Shen W, Wang K, Liu J, Xu B, Zhao J, Zhao C. 2012. Foxg1 has an essential role in postnatal development of the dentate gyrus. J Neurosci 32:2931–2949. doi:10.1523/JNEUROSCI.5240-11.2012

Villar-Cervino V, Molano-Mazon M, Catchpole T, Valdeolmillos M, Henkemeyer M, Martinez LM, Borrell V, Marin O. 2013. Contact repulsion controls the dispersion and final distribution of Cajal-Retzius cells. Neuron 77:457–471. doi:10.1016/j.neuron.2012.11.023

Weiss KH, Johanssen C, Tielsch A, Herz J, Deller T, Frotscher M, Forster E. 2003. Malformation of the radial glial scaffold in the dentate gyrus of reeler mice, scrambler mice, and ApoER2/VLDLR-deficient mice. J Comp Neurol 460:56–65. doi:10.1002/cne.10644

Yang A, Walker N, Bronson R, Kaghad M, Oosterwegel M, Bonnin J, Vagner C, Bonnet H, Dikkes P, Sharpe A, McKeon F, Caput D. 2000. p73-deficient mice have neurological, pheromonal and inflammatory defects but lack spontaneous tumours. Nature 404:99–103. doi:10.1038/35003607

Yoshida M, Assimacopoulos S, Jones KR, Grove EA. 2006. Massive loss of Cajal-Retzius cells does not disrupt neocortical layer order. Development 133:537–545. doi:10.1242/dev.02209

Zhao S, Chai X, Forster E, Frotscher M. 2004. Reelin is a positional signal for the lamination of dentate granule cells. Development 131:5117–5125. doi:10.1242/dev.01387

Zhao S, Frotscher M. 2010. Go or stop? Divergent roles of Reelin in radial neuronal migration. Neuroscientist 16:421–434. doi:10.1177/1073858410367521

