## Supplemental Figures for "Reelin Mediates Hippocampal Cajal-Retzius Cell Positioning and Infrapyramidal Blade Morphogenesis"

**SUPPORTING FIGURE LEGEND**

**Supporting figure 1.** Cell Death Analysis. Activated caspase-3 (green) staining at P4 showed no abnormal increase in cell death in *Reln* $^{CTRdel}$ hippocampus. Nuclei were stained using Hoechst (blue).

**Supporting figure 2.** Enlarged images of panels selected from Figure 5 are shown. **A:** The right color panel of Figure 5A. Scale bars, 100 μm. **B:** The right color panel of Figure 5B. Scale bars, 100 μm. **C:** The right panel of Figure 5D. Arrows indicate cells with elongated shape at the FDJ. Scale bars, 50 μm. **D:** The right color panel of Figure 5E. Scale bars, 100 μm.

**Supporting figure 3.** A comparison of the infrapyramidal blade (IPB) malformation between *Reln*$^{CTRdel}$ and *Apoer2*$^{null}$ mutants. *Apoer2*$^{null}$ mutant dentate gyrus also has a truncated IPB although not as severe as *Reln*$^{CTRdel}$. This phenotype appears more pronounced in the posterior sections; Images of the entire hippocampal region are displayed to show that cross-sections of the similar region were compared. Scale bars, 500 μm.

**Supporting figure 1**

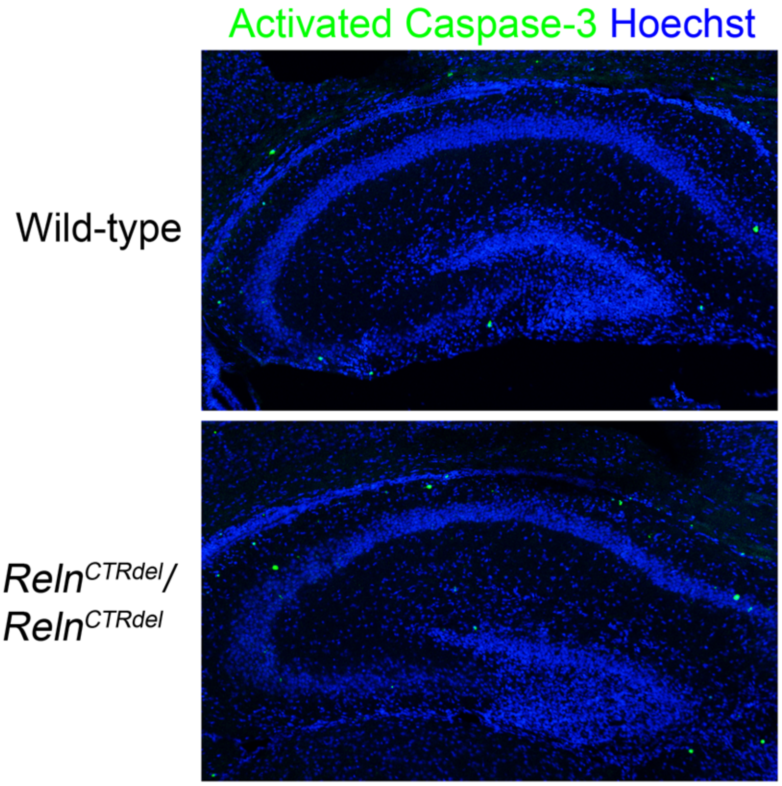

**Supporting figure 2**

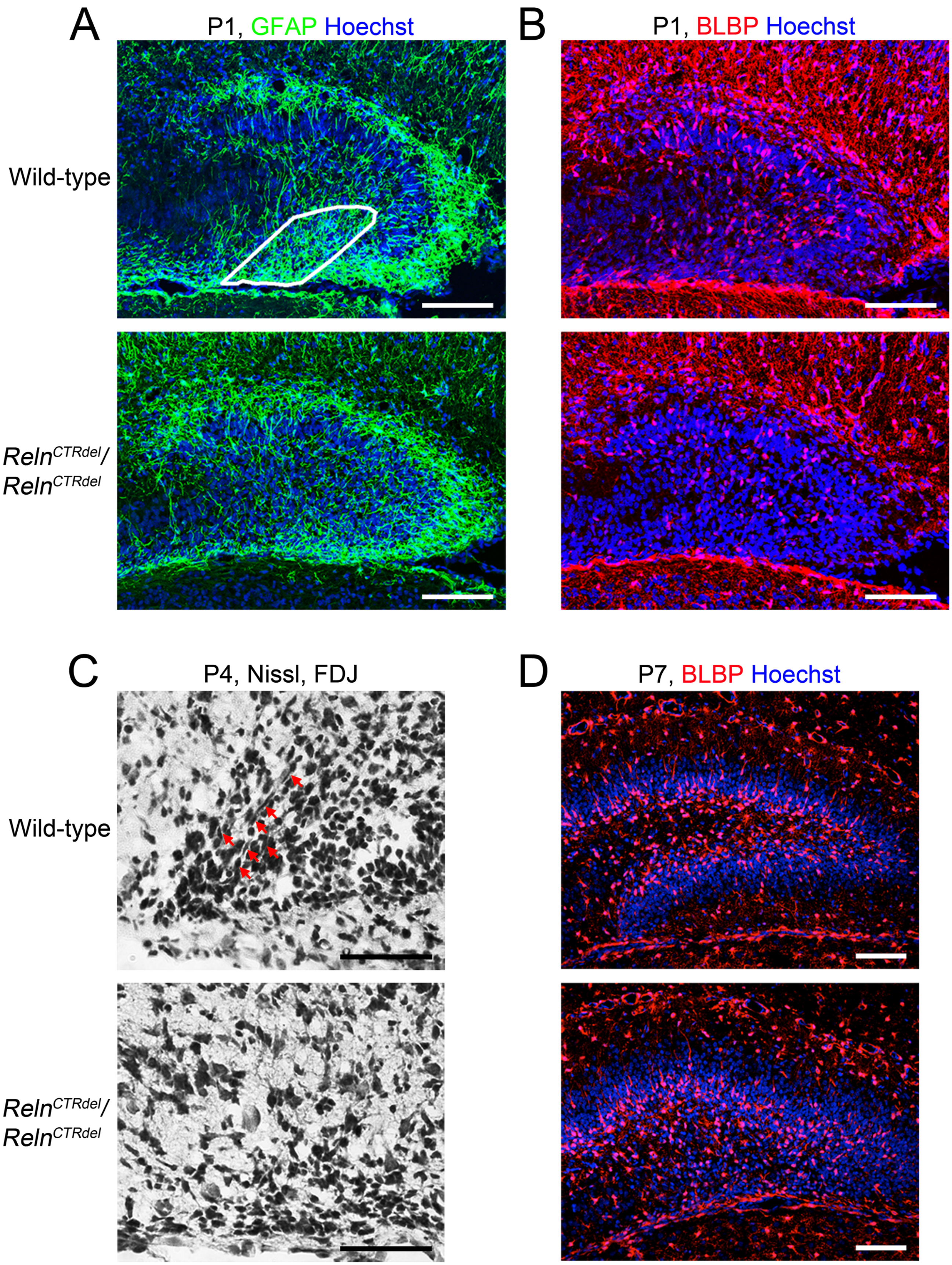

**Supporting figure 3**

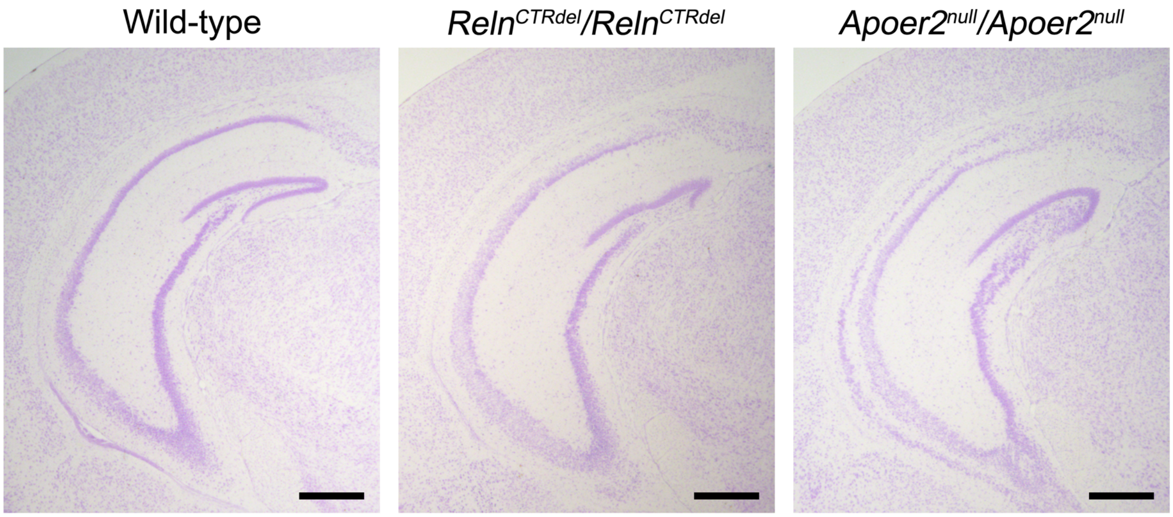
